# COMPUTATIONAL EVIDENCE FOR MULTI-LAYER CROSSTALK BETWEEN THE CADHERIN-11 AND PDGFR PATHWAYS

**DOI:** 10.1101/2022.11.08.514892

**Authors:** Zeynep Karagöz, Fiona R. Passanha, Lars Robeerst, Martijn van Griensven, Vanessa L. S. LaPointe, Aurélie Carlier

## Abstract

Various cell surface receptors play an important role in the differentiation and self-renewal of human mesenchymal stem cells (hMSCs). One example of such receptors are the cadherins, which maintain cell–cell adhesion and mechanically couple cells together. Recently, cadherin-11, which is a member of the type II classical cadherin family, has been shown to be involved in the fate commitment of hMSCs. Interestingly, cadherin-11 has no known intrinsic signaling activity and is thought to affect cell behavior via interactions with other cell surface receptors. Members of the platelet-derived growth factor receptor (PDGFR) family are hypothesized to be one of the interaction partners of cadherin-11. Experiments confirmed that PDGFR-α binding to extracellular cadherin-11 regions increases the PDGFR-α activity, whereas the interaction between PDGFR-β and cadherin-11 suppresses the activity of the growth factor receptor. Cadherin-11 knockdown experiments also decreased cell proliferation. These interactions between cadherin-11 and PDGFRs indicate a crosstalk between these receptors and their downstream signaling activities but the nature of this crosstalk is not entirely known. In this study, we used a computational model to represent the experimentally proven interactions between cadherin-11 and the two PDGFRs and we inspected whether the crosstalk also exists downstream of the signaling initiated by the two receptor families. The computational framework allowed us to monitor the relative activity levels of each protein in the network. We performed model simulations to mimic the conditions of previous cadherin-11 knockdown experiments and to predict the effect of crosstalk on cell proliferation. Overall, our predictions suggest the existence of another layer of crosstalk, namely between β-catenin (downstream to cadherin-11) and an ERK inhibitor protein (e.g. DUSP1), different than the crosstalk at the receptor level between cadherin-11 and PDGFR-α and -β. By investigating the multi-level crosstalk between cadherin and PDGFRs computationally, this study contributes to an improved understanding of the effect of cell surface receptors on hMSCs proliferation.

## INTRODUCTION

For decades we have known that signaling does not occur linearly, but through a complex network of interacting signals and pathways made up of signaling molecules (Barabási and Oltvai, 2004). Using wet laboratory experiments, a myriad of these signaling molecules have been identified and scientists have also tried to understand the crosstalk between them. However, these experiments have their limitations when it comes to studying the relationship between large networks of signaling pathways, as they can only isolate parts of the pathways and cannot look at the whole network. Computational models are better equipped to predict and analyze pathway crosstalk, as they offer a systematic way to conduct multivariate experiments that are impossible to perform *in vitro*, and they can also generate experimentally testable predictions. In our study, we have taken a wet laboratory experiment that studied the interaction between receptor tyrosine kinase, a cell surface receptor, and a specific cadherin, a cell adhesion protein (Takeichi, 2018) and we used computational modeling to add to the evidence and better understand the extent of the crosstalk.

There has been recent interest in the physical interaction between cadherin-11 and the two receptor tyrosine kinases (RTKs), PDGFR-α (Madarampalli et al., 2019;) and PGDGFR-β (Liu et al., 2020; Passanha et al., 2022) in fibroblasts and human mesenchymal stem cells (hMSCs) respectively. Passanha et al., 2022 reported that by using gene knockdown to temporarily decrease the expression of cadherin-11, the cadherin-11 knockdown cells have a more prolonged expression of phosphorylated ERK in the nuclei and these cells also show decreased proliferation. Similarly, Liu et al., 2019 showed that knocking down cadherin-11 also results in a decrease in proliferation. ERK is known to be downstream of the various RTKs including PGDGFR-β and so the current hypothesis is that the physical interactions between PGDGFR-β and cadherin-11 point towards a crosstalk between these two pathways. Although the receptor level interactions between RTKs and cadherins have been confirmed by wet laboratory experiments, we still know very little about the extent of this crosstalk as we lack experimental tools to investigate it. Here we want to use computational modeling to explore potential downstream interactions on top of the known receptor level interactions and expand the knowledge of this interesting crosstalk.

The ERK pathway is central to the progression of the cell cycle, proliferation, and growth of eukaryotic cells. It is known to be regulated by many growth factor receptors and thus part of many different signaling pathways, including the RTK and the cadherin pathways (Ramos, 2008). Using computational modeling, we isolated the RTK and cadherin-11 pathways and looked at how the crosstalk between these two pathways affects the ERK nuclear translocation leading to changes in cell proliferation. We observed that the downstream signaling in our model did not reflect the experimental evidence without a player between β-catenin (a subunit of the cadherin protein complex) and ERK that can influence proliferation. We, therefore, concluded that crosstalk at the receptor level between the RTK and cadherin-11 pathways alone is insufficient for cadherin-11 to influence hMSC proliferation through the ERK pathway, and that an additional level of crosstalk could be in place. By being able to study this interconnectedness between pathways, we have shown that our model can be used to describe the nature of the crosstalk between signaling molecules which is not always possible experimentally.

## METHODS

### Model Development

We built the signaling network in Figure 1 to include the PDGFR-α and PDGFR-β–induced ERK pathway as well as the cell–cell contact signaling via cadherin-11. The network represents the interactions of two adjacent cells and focuses on the intracellular response of one of these cells in contact. The receptor level interactions capture the experimentally established activation of PDGFR-α by cadherin-11 of the neighboring cell and the inhibition of PDGFR-β by cadherin-11 on the cell membrane. PDGFR-α and -β are activated by their ligands PDGF-α (a in Figure 1) and PDGF-β (b in Figure 1). The ERK pathway is activated downstream to the growth factor receptors, and follows the classical RAS-RAF-MEK-ERK cascade. ERK then activates its own inhibitor, DUSP1, which activates the cell cycle protein cyclin-D1. Downstream to cadherin-11, we included β-catenin that is inhibited by cadherin-11. B-catenin has shown to inhibit DUSP family proteins (Zeller et al., 2012), to interfere with the RAS-RAF-MEK-ERK cascade. Within this network, we also propose the inhibition of DUSP1 by β-catenin to be a key crosstalk mechanism, besides the experimentally established receptor-level interactions described earlier.

**Figure 1:**
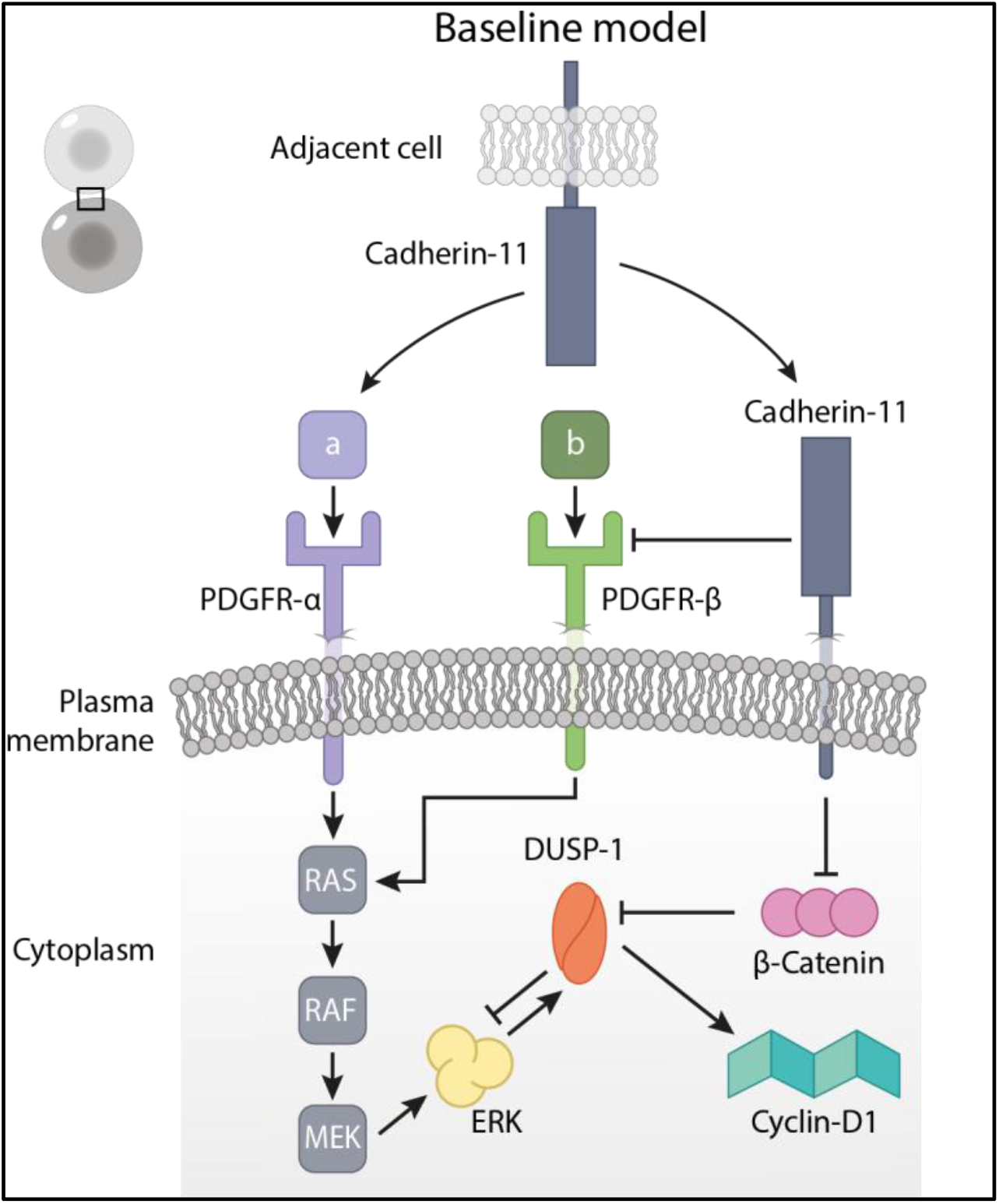
The schematic representation of receptor level crosstalk between growth factor receptors (PDGFR-α and PDGFR-β) and cadherin-11, and the proposed crosstalk between β-catenin and ERK via the ERK inhibitor DUSP1. Arrows represent activation and blunt arrows represent inhibition. The network represents the interactions of two adjacent cells (top left corner) and focuses on the intracellular response of one of these cells in contact.

We developed an ordinary differential equation (ODE) model to represent the known and suggested crosstalk between PDGFR and cadherin-11 as well as their downstream effectors. The ODEs 1–10 (Table 1) have the form suggested by Mendoza and Xenarios (Mendoza and Xenarios, 2006) to capture the qualitative behavior of cadherin-11 and PDGFR and their effect downstream, observed in the experiments by both Madarampalli et al., 2019 and Passanha et al., 2022. We chose this type of equation as it allows a signaling network to be translated into a continuous dynamical system and study its stable steady state and qualitative behavior without the need for precise data on the signaling stoichiometry and kinetics.

**Table 1:** Ordinary differential equations governing the activity levels of each protein in the model network.

| Nr. | Equation |
| --- | --- |
| 1 | $\frac{dcdh11}{dt} = \frac{-e^{0.5 \times h} + e^{-h \times (\omega_{cdh11}-0.5)}}{(1 - e^{0.5 \times h}) \times (1 + e^{-h \times (\omega_{cdh11}-0.5)})} - (k_i \times cdh11))$ |
| 2 | $\frac{dpdgfrA}{dt} = \frac{-e^{0.5 \times h} + e^{-h \times (\omega_{pdgfrA}-0.5)}}{(1 - e^{0.5 \times h}) \times (1 + e^{-h \times (\omega_{pdgfrA}-0.5)})} - (k_i \times pdgfrA))$ |
| 3 | $\frac{dpdgfrB}{dt} = \frac{-e^{0.5 \times h} + e^{-h \times (\omega_{pdgfrB}-0.5)}}{(1 - e^{0.5 \times h}) \times (1 + e^{-h \times (\omega_{pdgfrB}-0.5)})} - (k_i \times pdgfrB))$ |
| 4 | $\frac{dras}{dt} = \frac{-e^{0.5 \times h} + e^{-h \times (\omega_{ras}-0.5)}}{(1 - e^{0.5 \times h}) \times (1 + e^{-h \times (\omega_{ras}-0.5)})} - (k_i \times ras))$ |
| 5 | $\frac{draf}{dt} = \frac{-e^{0.5 \times h} + e^{-h \times (\omega_{raf}-0.5)}}{(1 - e^{0.5 \times h}) \times (1 + e^{-h \times (\omega_{raf}-0.5)})} - (k_i \times raf))$ |
| 6 | $\frac{dmek}{dt} = \frac{-e^{0.5 \times h} + e^{-h \times (\omega_{mek}-0.5)}}{(1 - e^{0.5 \times h}) \times (1 + e^{-h \times (\omega_{mek}-0.5)})} - (k_i \times mek))$ |
| 7 | $\frac{derk}{dt} = \frac{-e^{0.5 \times h} + e^{-h \times (\omega_{erk}-0.5)}}{(1 - e^{0.5 \times h}) \times (1 + e^{-h \times (\omega_{erk}-0.5)})} - (k_i \times erk))$ |
| 8 | $\frac{ddusp1}{dt} = \frac{-e^{0.5 \times h} + e^{-h \times (\omega_{dusp1}-0.5)}}{(1 - e^{0.5 \times h}) \times (1 + e^{-h \times (\omega_{dusp1}-0.5)})} - (k_i \times dusp1))$ |
| 9 | $\frac{dBcat}{dt} = \frac{-e^{0.5 \times h} + e^{-h \times (\omega_{Bcat}-0.5)}}{(1 - e^{0.5 \times h}) \times (1 + e^{-h \times (\omega_{Bcat}-0.5)})} - (k_{iB} \times Bcat))$ |
| 10 | $\frac{dcyclinD1}{dt} = \frac{-e^{0.5 \times h} + e^{-h \times (\omega_{cyclinD1}-0.5)}}{(1 - e^{0.5 \times h}) \times (1 + e^{-h \times (\omega_{cyclinD1}-0.5)})} - (k_i \times cyclinD1))$ |

The model equations (Table 1) represent the rate of change in the activity level of each protein in the network. Here the “activity level” indicates the net effect of a protein in its active form (either in the phosphorylated form or otherwise functional). For example, when we mention “activity of ERK” in relation to this model, we mean “phosphorylated ERK in the nucleus, where it is active”. The activity level of proteins varies between 0 and 1 (1 being maximum activity) and it is unitless, due to the nature of the ODEs described in Mendoza and Xenarios, 2006. Each ODE has an activation term and a decay term. The decay term mimics the autoinhibition or the inactivation of proteins in the cell over time. The general decay parameter, *k*_*i*_, was set to 1 for all proteins for simplicity. Only for the β-catenin decay parameter, we used *k*_*iB*_ = 2 to compensate for its high initial activity and to account for the involvement of β-catenin in other intracellular pathways (Valenta et al., 2012). The activation term in the ODEs includes a parameter omega (*ω*) which is specific to each protein and the values of *ω* can be calculated using the parameters in Table 2. *ω* represents the total input to the activity of each protein at a particular time. The alpha (α) parameters represent activation and the beta (β) parameters represent inhibition between proteins. For example, PDGFR-β is activated by its own ligand PDGF-β while it is also inhibited by cadherin-11 therefore we use 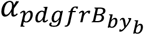 to represent the activation and 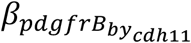 to represent the inhibition in *ω*_*pdgfrB*_. As such, using the parameters in Table 2, we mathematically built the network given in Figure 1.

**Table 2:**
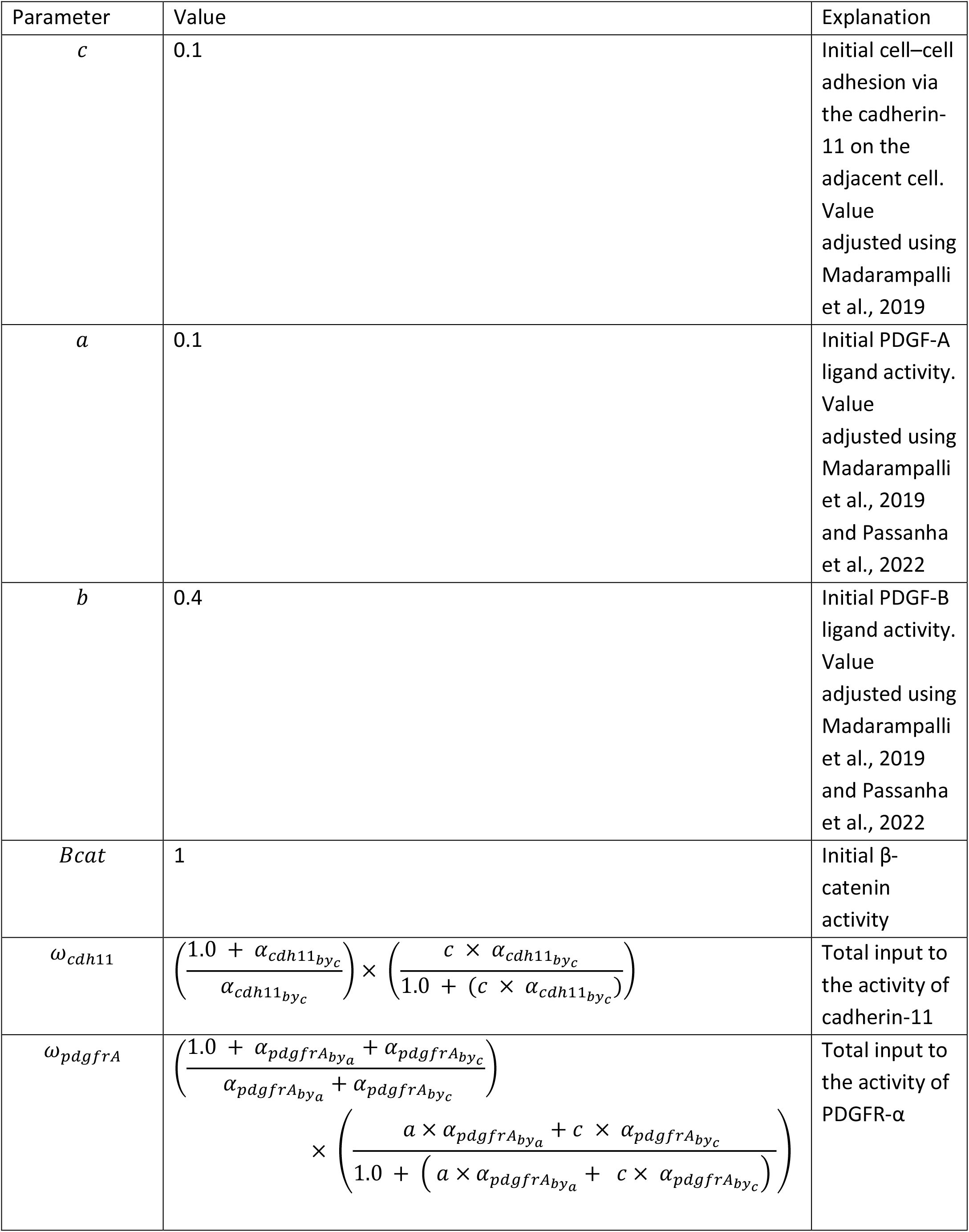
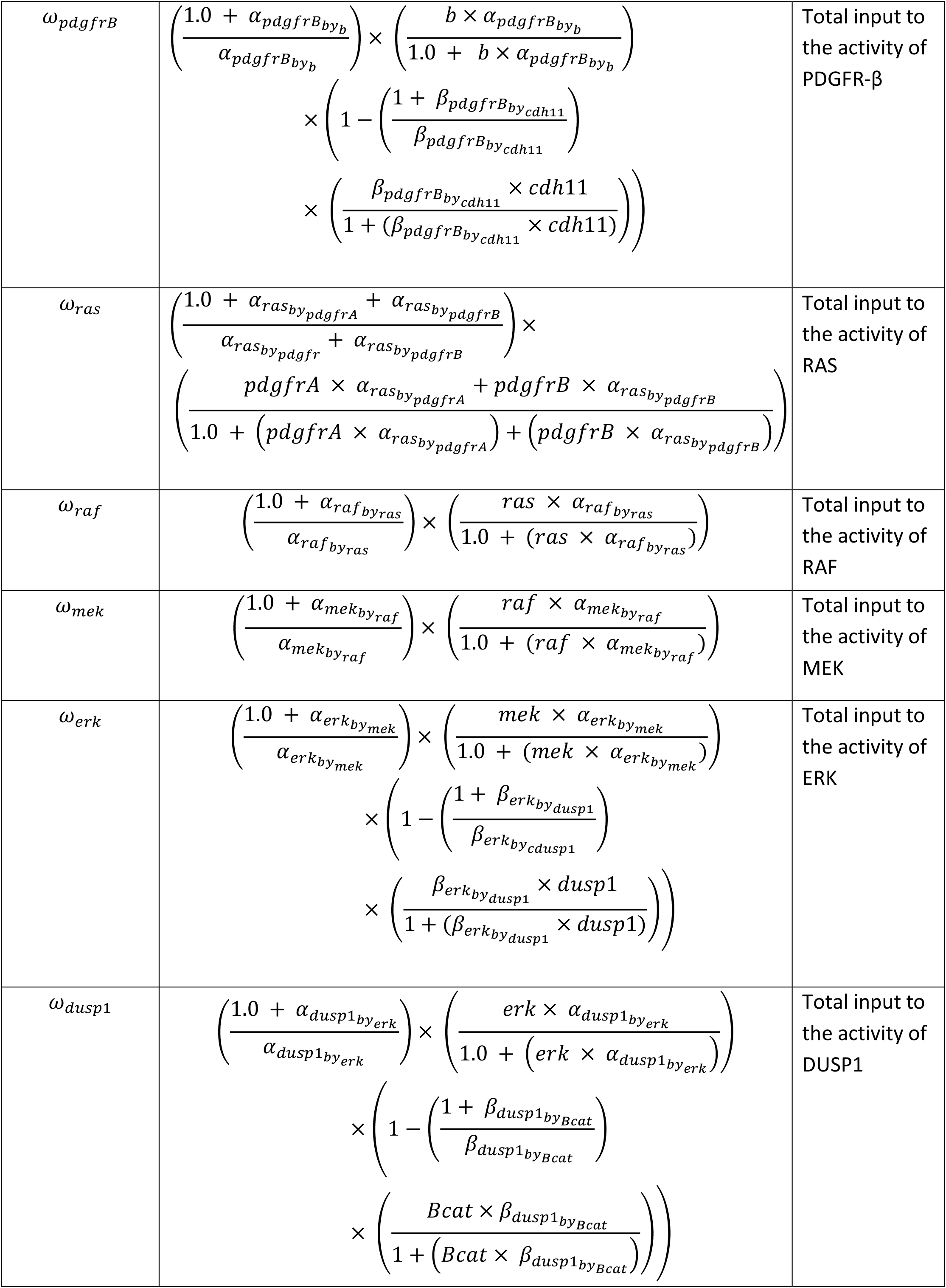
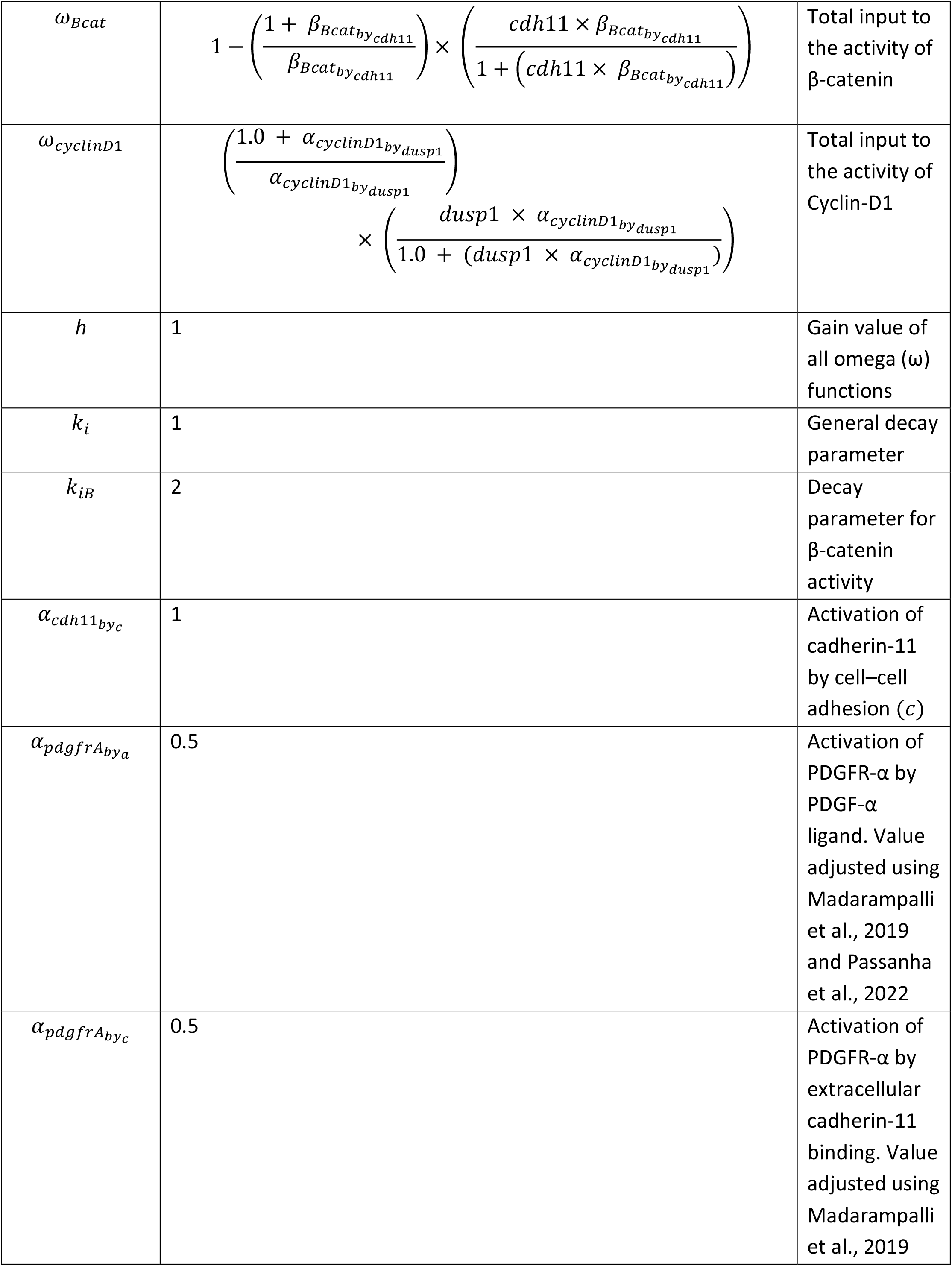
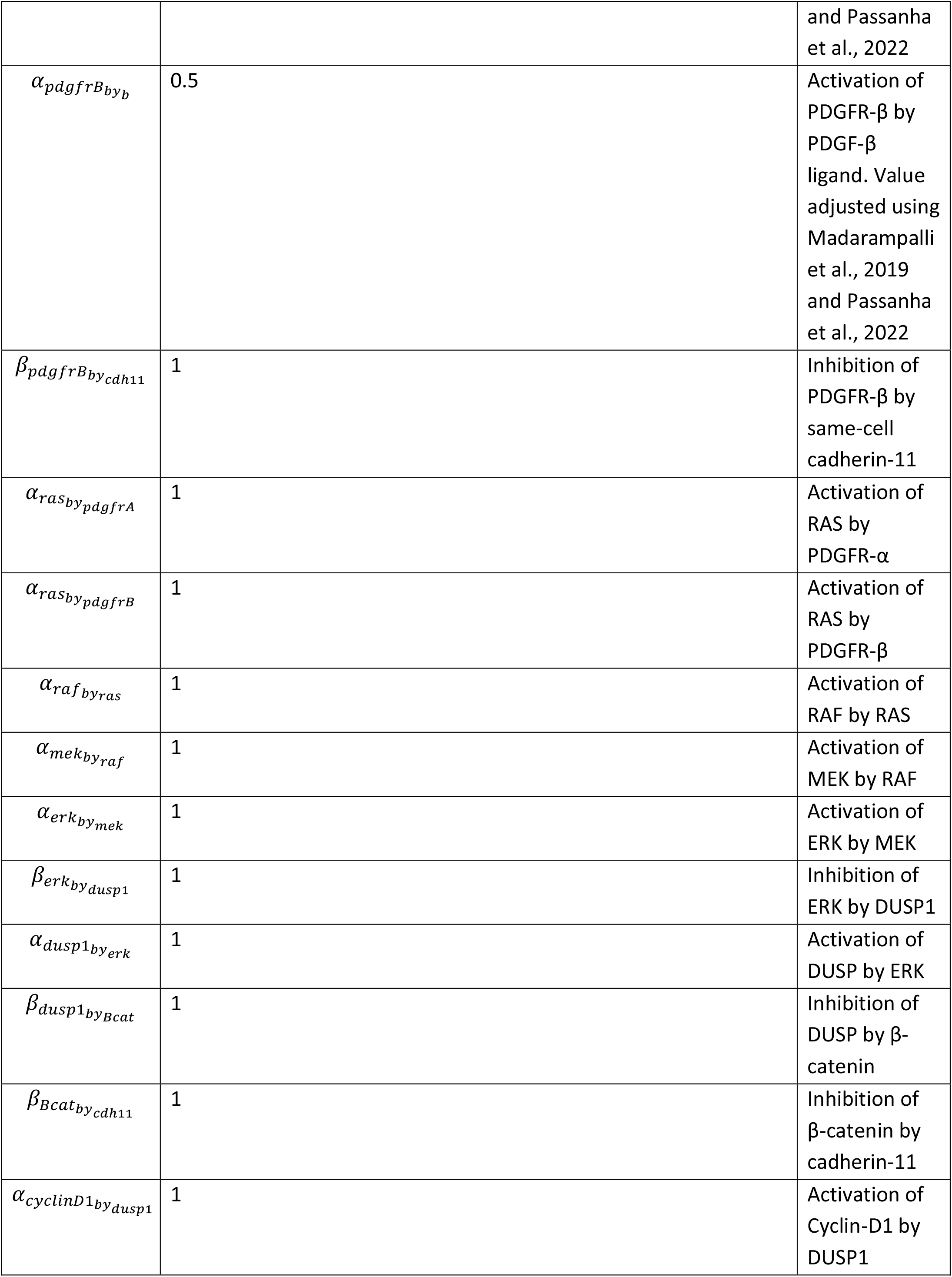
Parameter values used in the baseline simulation.

The parameters *a* and *b* and the initial activity levels of PDGF-A and PDGF-B were adjusted to meet the quantitative measurements of PDGFR-α and PDGFR-β activity levels in cadherin-11 knockdown experiments by Madarampalli et al., 2019 and Passanha et al., 2022, respectively. The remaining parameters were set to a default value of 1, as suggested by Mendoza and Xenarios, 2006 in case of insufficient experimental data. It is important to note that by using this type of ODEs, we were able to capture the qualitative behavior of the whole network and the changes in the activity levels of each protein at the steady state, while the time, and consequently also the dynamics, were arbitrary, as described in detail in Mendoza and Xenarios, 2006.

### Simulations

We used the Virtual Cell software (VCell) version 7.4.0 (Schaff et al., 1997; Cowan et al., 2012) to simulate the network. The baseline model, all different simulation setups, and the results can be accessed within the VCell software (Access to the VCell model will be granted upon request during the preprint period. Once the paper is peer-reviewed, the VCell model will be publicy available).

We first performed a simulation of the baseline model (Table 3: Baseline model) using the parameters in Table 2. This simulation provided a baseline steady state activity for each protein in the network. In the next simulation (Table 3: Cadherin-11 knockdown) we set the initial cadherin-11 activity to zero on both the cell for which the intracellular signaling is modeled and on the adjacent cell, mimicking the experimental cadherin-11 knockdown. We then compared the steady state activity levels of the cadherin-11 knockdown simulation to the baseline activity levels. Other simulations have been performed to test the effect of different crosstalk modes in the network, namely the crosstalk only at the receptor level, crosstalk only downstream to the receptors or both at the same time. These simulations have been explained in the text where relevant, and the corresponding parameter changes are given in Table 3.

**Table 3:**
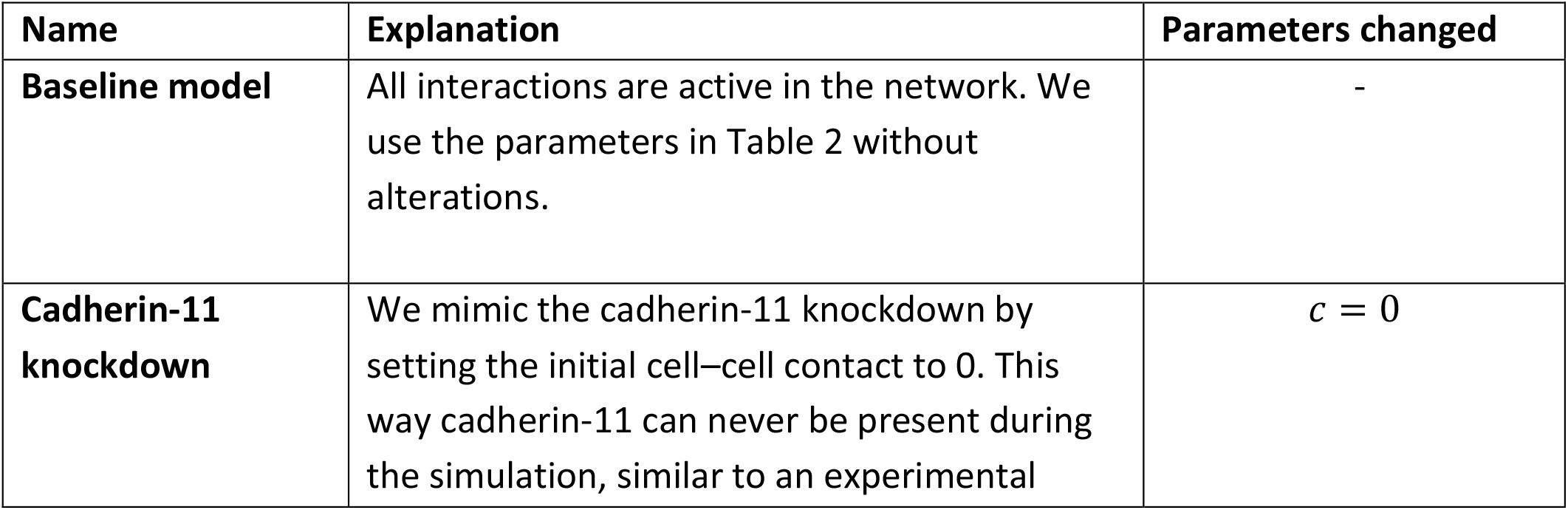
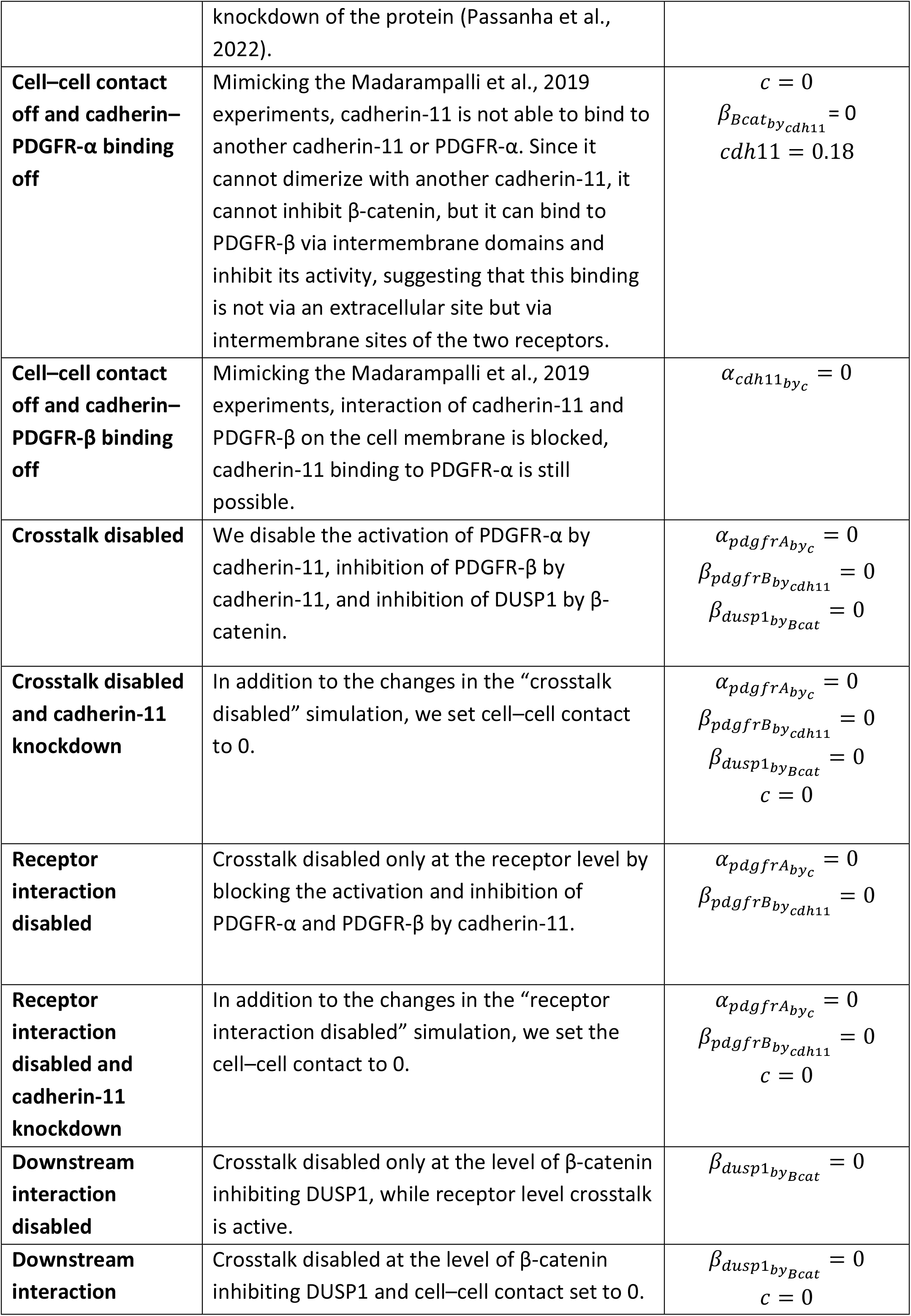
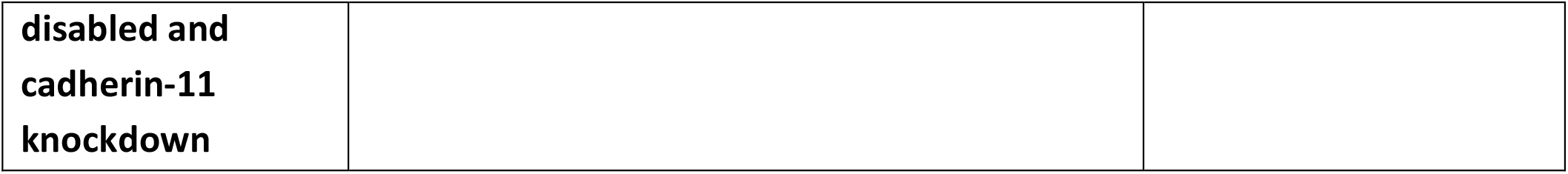
Setups of the four main simulations referred to in the main figures.

### Parameter Scan

To ensure our choice of parameters around the proposed crosstalk between DUSP1 and β-catenin did not force the system to behave in a biased way, we performed a parameter scan in groups of two at a time. First, different values (0.1, 0.5, 1.0, 5.0, 7.0, 10.0) of the parameter determining the inhibition of DUSP1 in the network 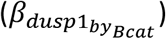 were scanned against different values of *Bcat* (0.1, 0.3, 0.5, 0.7, 1.0) and 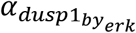 0.1, 0.5, 1.0, 5.0, 7.0, 10.0), which are the two parameters that contribute to the activation of DUSP1 in the network. Second, the parameter that contributes to the inhibition of ERK by DUSP1 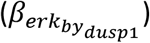 was scanned against the parameter that contributes to the activation of ERK 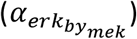. Last, 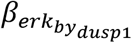 was scanned against the parameter that contributes to the activation of cyclin-D1 by DUSP1 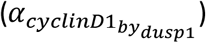. For the last two scans, we used the same parameter space for all parameters (0.1, 0.5, 1.0, 5.0, 7.0, 10.0). For all four parameter scans, parameters were varied simultaneously to cover all 25 combinations of the parameter sets.

As the output of the parameter scan, we reported the difference in steady state activity levels of cyclin-D1 and ERK as a measure of the change in cell proliferation compared to the baseline. Simulation with the baseline model parameters (all equal to 1) resulted in equal activity levels for cyclin-D1 and ERK. If for a parameter set, the cyclin-D1 activity was higher than ERK activity we classified this as “increased proliferation” compared to the baseline. If the cyclin-D1 activity was lower than ERK activity, we classified this as “decreased proliferation” compared to the baseline. For the cyclin-D1 activity matching the ERK activity we classified this as no change in proliferation compared to the baseline model parameter set. The results of this analysis are summarized in Figure 3 and the numerical results are given in Tables S1-4.

## RESULTS

First, in order to ensure that the baseline model captured the experimentally established activation of PDGFR-α by cadherin-11 on an adjacent cell and the inhibition of PDGFR-β by cadherin-11 in the cell membrane, we ran the baseline model simulation as is (Table 3: Baseline model), the cadherin-11 knockdown simulation (Table 3: Cadherin-11 knockdown), a simulation where the receptor interaction was disrupted via blocking the cadherin-11 binding to PDGFR-α (Table 3: Cell–cell contact off and cadherin–PDGFR-α binding off), and a simulation where the receptor interaction was disrupted by blocking the cadherin-11 binding to PDGFR-β (Table 3: Cell–cell contact off and cadherin–PDGFR-β binding off). In line with Passanha et al., 2022, the cadherin-11 knockdown resulted in a 50% decrease in PDGFR-α and a 30% increase in PDGFR-β activity compared to the baseline simulation (Figure S1, Cadherin-11 knockdown). Also in line with Madarampalli et al., 2019, the absence of cadherin-11 binding to PDGFR-α resulted in 50% lower PDGFR-α activity while PDGFR-β activity did not change compared to the baseline simulation (Figure S1, Cell–cell contact off and cadherin–PDGFR-α binding off). Lastly, the absence of the cadherin-11 binding to PDGFR-β only increased the PDGFR-β activity by 30% but did not affect the PDGFR-α activity compared to the baseline simulation (Figure S1, Cell–cell contact off and cadherin–PDGFR-β binding off). It is important to note that these simulations were done only to confirm that the parameters of the model had been adjusted correctly for the part of the model network for which we have experimental evidence. As such, these results are not providing proof for the whole network, but they are important in linking the model to prior experiments.

Next, we explored the remaining parts of the model in a cadherin-11 knockdown simulation (Table 3: Cadherin-11 knockdown). Figure 2 summarizes the changes in activity levels of the proteins in the network in a cadherin-11 knockdown simulation compared to their baseline activity levels. The two proteins whose activities were inhibited by cadherin-11 in the model, namely β-catenin and PDGFR-β, showed an overall increase in activity, as expected (Figure 2A, Figure 2B Cadherin-11 knockdown). PDGFR-α, on the other hand, had lower activity in the cadherin-11 knockdown, as it is normally activated by cadherin-11, alongside its own ligand (Figure 2A, Figure 2B Cadherin-11 knockdown). The net effect of the cadherin-11 knockdown on ERK was an increase in activity, while both DUSP1 and cyclin-D1 showed decreased activities (Figure 2A, Figure 2B, Cadherin-11 knockdown). In the context of our computational model, we interpreted the decreased cyclin-D1 activity compared to the baseline in the cadherin-11 knockdown as a decrease in cell proliferation.

**Figure 2:**
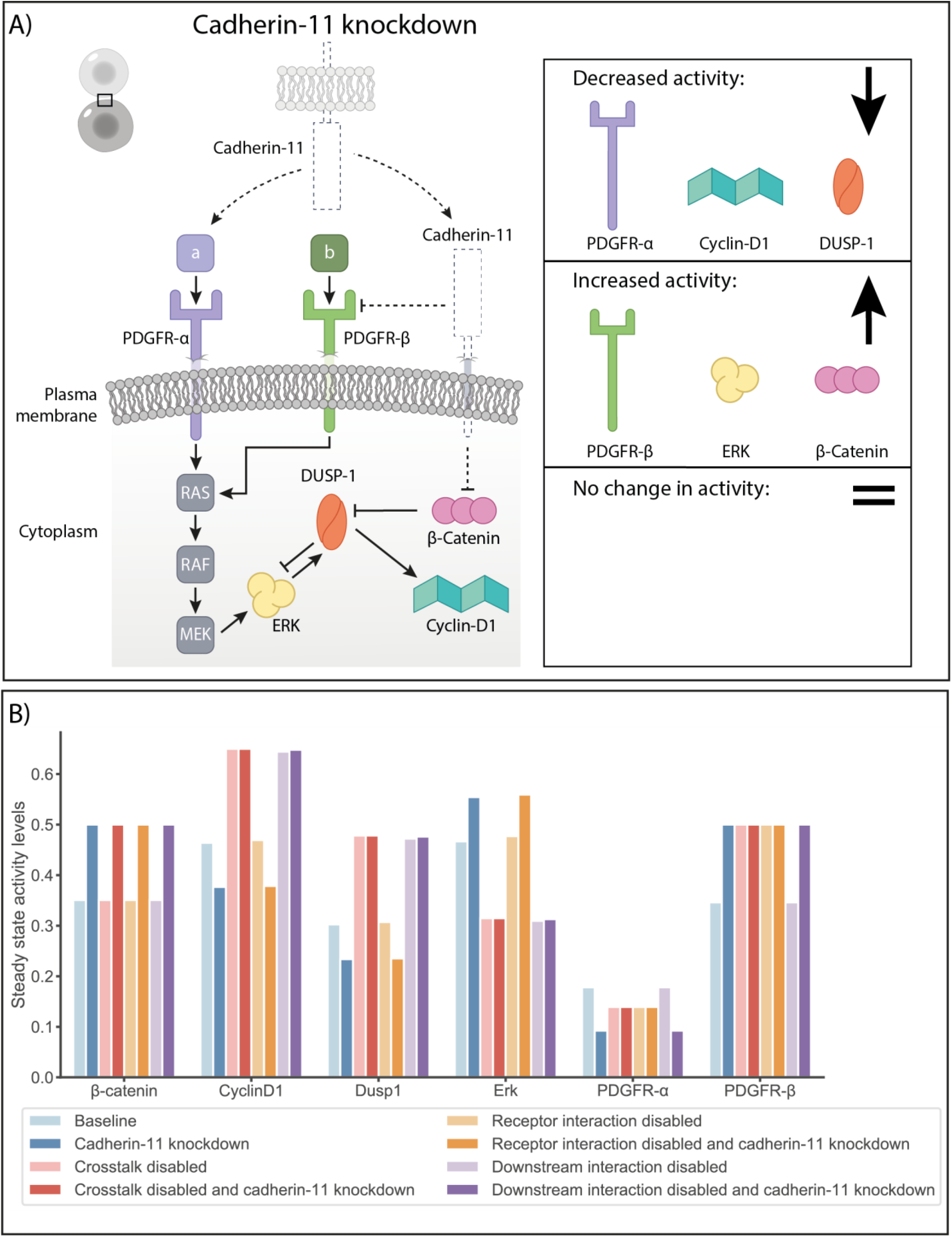
**A)** Cadherin 11 knockdown simulation results compared to the baseline model simulation: The dashed lines indicate where the model components were modified in the simulation setup. The steady state activity levels of PDGFR-α, cyclin-D1 and DUSP1 decreased, while the activity levels of PDGFR-β, ERK and β-catenin increased in the cadherin-11 knockdown compared to the baseline simulation. The decrease in PDGFR-α activity and the increase in PDGFR-β activity agree with the experimental results (Madarampalli et al., 2019; Passanha et al., 2022). The proposed crosstalk protein, DUSP1, had decreased activity which resulted in increased ERK activity and decreased cyclin-D1 activity, which we interpreted as decreased proliferation. **B)** Summary of steady state activity levels of proteins in the network under different simulation setups. We compared the steady state activity levels in the cadherin-11 knockdown for each different setup (baseline, crosstalk disabled, receptor interaction disabled, and downstream interaction disabled).

Having observed that the cadherin-11 knockdown simulation of the baseline model setup reflected the experimental observations of increased ERK activity and decreased proliferation in hMSCs, we moved to testing the contribution of different modes of crosstalk to these results. When the crosstalk at the receptor level (i.e. activation of PDGFR-α by cadherin-11 on an adjacent cell membrane and inhibition of PDGFR-β by cadherin-11 on the cell’s own membrane) was disabled and the cadherin-11 knockdown simulation was repeated (Table 3: Receptor interaction disabled and cadherin-11 knockdown), the same ERK and cyclin-D1 activities were reached as in the cadherin-11 knockdown simulation using the baseline model (Figure 2B, Receptor interaction disabled and cadherin-11 knockdown). In other words, the removal of receptor level interactions did not affect the cadherin-11 influence on proliferation. This implies the receptor level crosstalk is not the primary mode of crosstalk that maintains the cadherin-11–dependent cell proliferation.

In order to test the effect of the crosstalk at the level of β-catenin and DUSP1, we modified the model to remove both the inhibition of DUSP1 by β-catenin and the receptor crosstalk (Table 3: Crosstalk disabled). In this case, β-catenin activity remained the same as in the baseline simulation, while PDGFR-α had a lower activity due to the lack of activation by cadherin-11 (Figure 2B, Crosstalk disabled versus Baseline). On the contrary, PDGFR-β activity increased in the absence of crosstalk, as it was no longer inhibited by cadherin-11. The net effect of not having the proposed multi-layered crosstalk between the two pathways resulted in decreased ERK activity and increased DUSP1 and cyclin-D1 activity (i.e., proliferation) compared to the baseline simulation (Figure 2B, Crosstalk disabled versus Baseline). When we simulated a cadherin-11 knockdown case for this version of the model (Table 3: Crosstalk disabled and cadherin-11 knockdown), we observed that the cadherin-11–mediated cell proliferation was disrupted. Unlike in the cadherin-11 knockdown, where the inhibition of DUSP1 by β-catenin was still active (Figure 2A, Figure 2B Cadherin-11 knockdown versus Baseline), we did not observe any change in the activities of PDGFR-α, PDGFR-β, Cyclin-D1, DUSP1 and ERK (Figure 2B, Crosstalk disabled and cadherin-11 knockdown versus Crosstalk disabled, and Figure S2). Only the activity of β-catenin was increased, which was due to the absence of inhibition by cadherin-11 (Figure 2B, Crosstalk disabled and cadherin-11 knockdown versus Crosstalk disabled, and Figure S2).

Next, we explored whether the crosstalk at the level of DUSP1 and β-catenin could maintain the cadherin-11–mediated cell proliferation on its own. To do this, we kept the receptor level crosstalk active (cadherin-11 activating PDGFR-α and inhibiting PDGFR-β), disabled the inhibition of DUSP1 by β-catenin (Table 3: Downstream interaction disabled), and performed a cadherin-11 knockdown simulation with this setup (Table 3: Downstream interaction disabled and cadherin-11 knockdown). We observed the same changes in protein activity levels with this setup as in the “crosstalk disabled” simulation (Figure 2B crosstalk disabled versus downstream interaction disabled). These results indicated that cadherin-11 could mediate cell proliferation via its downstream effector β-catenin by inhibiting DUSP1 which is an ERK inhibitor. According to our model predictions, receptor level interactions between cadherin-11 and PGDFR-α and PDGFR-β, on the other hand, were insufficient to orchestrate the cadherin-11–mediated cell proliferation.

We also explored the parameters related to the proposed crosstalk at the level of β-catenin and DUSP1 with a parameter scan (Figure 3). This enabled us to ensure our choice of parameters in the baseline model was not forcing the network to behave in a biased way and to decide which protein–protein interactions had more weight in controlling the proliferative state of the network.

**Figure 3:**
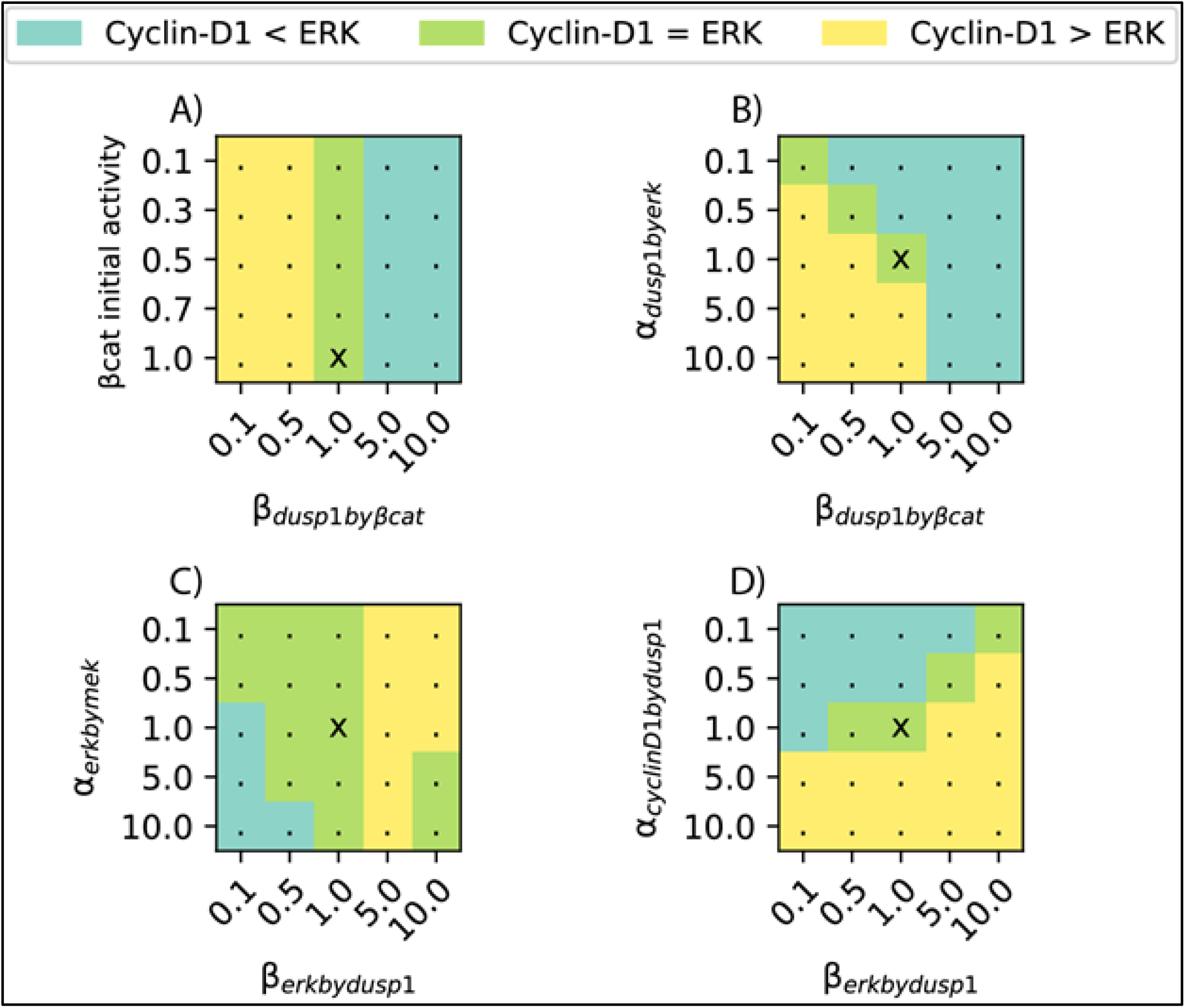
Change in proliferation status for varying parameters influencing the proposed crosstalk. The scan was performed using the numerical values on the axes for the respective parameters. Baseline parameter sets have been marked with an x in each plot. For each parameter set, the difference between the steady state activity level of cyclin D1 and ERK were compared. A) β-catenin initial activity and the inhibition of DUSP1 by β-catenin, B) activation of DUSP1 by ERK and inhibition of DUSP1 by β-catenin C) activation of ERK by MEK and inhibition of ERK by DUSP1 and D) activation of cyclin-D1 by DUSP1 and inhibition of ERK by DUSP1. Cyclin-D1 activity > ERK activity indicates increased proliferation compared to the baseline parameter set, cyclin-D1 activity = ERK activity indicates sustained proliferation as in the baseline parameter set, cyclin-D1 activity < ERK activity indicates decreased proliferation compared to the baseline parameter set.

When using the baseline model parameters, ERK and cyclin-D1 activities were equal at the steady state. In addition, for all parameter sets and in each parameter scan, there were multiple other parameter combinations that resulted in equal ERK and cyclin-D1 activity (Figure 3, green areas). This ensured the baseline model results were not unique and were therefore not resulting from any particular parameter settings.

The first scan investigated the parameters influencing the inhibition of DUSP1 by β-catenin. It revealed that the initial β-catenin activity level was not decisive on the proliferation status of the network when compared to the inhibition of DUSP1 by β-catenin 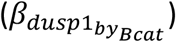 (Figure 3A). In other words, the activity of β-catenin itself did not affect the steady state of Cyclin-D1 and ERK, whereas they were affected by the strength at which β-catenin inhibited DUSP. For values of 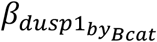 greater than 1, the ERK activity was higher than the cyclin-D1 activity (i.e., the proliferation was lower than in the baseline model). For values less than 1 of 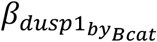, the Cyclin-D1 activity was higher than the ERK activity (Figure 3A).

The second scan was performed between 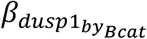 and 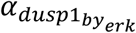, which are parameters that affect the strength of inhibition and activation of DUSP1, respectively. The results suggested that for the strong inhibition of DUSP1 by β-catenin 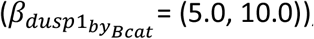, the network exhibited decreased proliferation (Figure 3B, Cyclin-D1 < ERK), irrespective of the value of DUSP1 activation by ERK 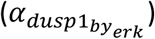. For the weaker inhibition of DUSP1 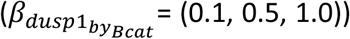, on the other hand, the increased value of DUSP1 activation by 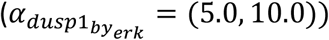 resulted in increased proliferation (Figure 3B, Cyclin-D1 > ERK). There exists a trade-off for values lower than or equal to 1 for both parameters, when the inhibition of DUSP1 by β-catenin exceeded the value of the activation of DUSP1 by ERK, the network exhibited decreased proliferation (Figure 3B, Cyclin-D1 < ERK). Whereas when 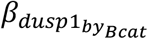 was smaller than 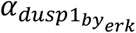 the proliferation increased (Figure 3B, Cyclin-D1 < ERK).

The third parameter set explored the activation and inhibition of ERK by the downstream effectors of the PDGFRs and DUSP1, respectively. The results suggested that, apart from the combination of the weakest possible inhibition of ERK by DUSP1 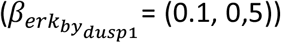 and the strongest possible activation of ERK by MEK 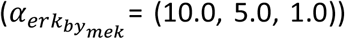, the network exhibited normal or increased proliferation (Figure 3C). In general, the stronger inhibition of ERK by DUSP1 resulted in increased proliferation, suggesting the control over ERK inhibition to be an important mechanism in controlling the proliferative status of the network.

In the final parameter scan, we further explored the interplay between the inhibition of ERK by DUSP1 and the activation of cyclin-D1 by DUSP1. For the stronger activation of cyclin-D1 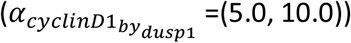, the proliferation was always higher than baseline, regardless of the strength of ERK inhibition by DUSP1 (Figure 3D). However, for weaker activation of cyclin-D1, the degree of ERK inhibition by DUSP1 could compensate and sustain the same level of proliferation or even increase it. For 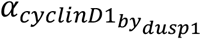 between 0.1 and 1.0, increasing 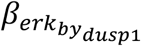 increased the proliferation (Figure 3D).

## DISCUSSION

There has been a recent increase in exploring the different (signaling) roles cadherins play in various cell types other than establishing cell to cell contact (Halbleib and Nelson, 2006; Niessen et al., 2011; Maître and Heisenberg, 2013). One of the many ways in which cadherins participate in various signaling roles is through their crosstalk with RTKs (Chiasson-Mackenzie and McClatchey, 2018). For example, there is mounting evidence of the crosstalk between cadherin-2 and the fibroblast growth factor receptors, as well as cadherin-1 and epidermal growth factor receptor (Andl and Rustgi, 2004; Nguyen and Mège, 2016). Likewise, recent studies have shown that cadherin-11 specifically interacts with the RTKs PDGFR-α (Madarampalli et al., 2019) and PDGFR-β (Liu et al., 2020; Passanha et al., 2022). Passanha et al., 2022 also reported that cadherin-11–knockdown hMSCs have more phosphorylated ERK-positive nuclei and show decreased proliferation. Similarly, Liu et al., 2020 have shown that knocking down cadherin-11 also results in decreased proliferation. Although these experimental observations point to crosstalk between PDGFR-α, PDGFR-β and cadherin-11 at the receptor level in hMSCs, crosstalk between the signaling proteins that function downstream to these receptors remains to be explored. This knowledge could improve our understanding of hMSC proliferation and fate commitment, both of which are central to regenerative medicine research.

Downstream to the PDGFRs is the MAPK/ERK pathway which contains multifunctional proteins. Downstream to cadherin-11 is β-catenin which is known to be involved in other cellular activities. The complexity and multifunctionality of the downstream signaling make it experimentally challenging to study the possible ways by which PDGFRs and cadherin-11 engage in crosstalk, other than at the receptor level. To address this challenge, we created a computational model of the cadherin-11 and PDGFR signaling network that qualitatively matches the experimental observations at the level of receptor interactions. With our model, we were able to provide *in silico* evidence for an additional layer of crosstalk (i.e., at the downstream level between β-catenin and an ERK inhibitor DUSP1), beyond the known crosstalk happening at the receptor level between PDGFRs and cadherin-11.

Computational modeling enabled us to isolate the RTK and cadherin-11 pathways from other cellular signaling pathways they interact with (e.g., integrins, G-protein coupled receptors (GPCRs), TGFβ, *etc*.) and explore the role of each mode of crosstalk (receptor level, downstream signaling and a combination of both) in the network in a way that is not possible experimentally. Disabling all possible crosstalk in the network, resulted in complete isolation of the ERK pathway from the cadherin-11, therefore the cadherin-11 knockdown did not affect cyclin-D1 or ERK activity (Figure 2B, crosstalk disabled and cadherin-11 knockdown compared to crosstalk disabled). Disabling the crosstalk only at the receptor level (i.e., between PDGFR-α and cadherin-11, and PDGFR-β and cadherin-11) and allowing the downstream interaction (i.e. β-catenin inhibiting DUSP1) rescued the activity level patterns and we observed that in case of a cadherin-11 knockdown while the receptor interaction was disabled, the ERK activity increased and cyclin-D1 activity decreased (Figure 2B, Receptor interaction disabled and cadherin-11 knockdown) matching the baseline simulation (Figure 2B, baseline and cadherin-11 knockdown). This suggests, the downstream interaction alone, can sustain the crosstalk between the cadherin-11 and PDGFR pathways and explain the observed decrease in proliferation in the cadherin-11 knockdown experiments. The observation that disabling the downstream crosstalk via β-catenin and the ERK inhibitor DUSP1 alone resulted in the same pattern as disabling all crosstalk (Figure 2B, Downstream interaction disabled versus crosstalk disabled) strengthens this hypothesis.

According to our simulations, the control over ERK inhibition in the network is very important. To ensure that our observations are not heavily influenced by our choice of parameters for the proposed crosstalk, we performed a sensitivity analysis. The results implied that for the strong inhibition of ERK (high 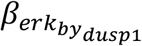 by DUSP1, regardless of the degree of activation of ERK by MEK 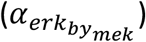, the system will have increased or at least sustained proliferation. Conversely, for strong ERK activation by MEK and weak inhibition of ERK by DUSP1, the proliferation was decreased (Figure 3).

Overall, the control over the inhibition of ERK seems to be critical in terms of changing the proliferative state of the system. This can be due to the fact that during cell proliferation, ERK is rapidly inactivated at the transition of the G1 to S phase (Meloche, 1995; Mebratu and Tesfaigzi, 2009). Prolonged ERK activation in the nucleus arrests the cell cycle at the G1 phase and therefore the cell does not undergo mitosis (Yamamoto et al., 2006). Therefore, the tight regulation of ERK (inhibiting its activity) is necessary for cell proliferation. We suggest cadherin-11 and its downstream effector β-catenin play a part in the control over ERK inhibition, explaining why the cadherin-11 knockdown results in decreased hMSC proliferation (as shown experimentally by Passanha et al., 2022).

In our model, we propose the ERK inhibitor in this network to be DUSP1 because the DUSP family of proteins dephosphorylate various members of the MAPK family including ERK (Huang and Tan, 2012; Chen et al., 2019). DUSP1 is a nuclear phosphatase that binds to ERK which leads to its dephosphorylation or inactivation, and reciprocally, ERK also promotes the activity of DUSP1 (Ferguson et al., 2016). This is not to say that other members of the DUSP family are not involved, but we chose to only highlight DUSP1 in our model for simplicity. We note that other signaling molecules can also regulate the activity of DUSPs, representing an additional signaling component that could be added to the model in the future. The DUSP family of proteins has also been shown to interact with β-catenin to interfere with the MAPK signaling cascade in murine liver cells (Zeller et al., 2012). Therefore, we suggest that in the scope of the RTK and cadherin-11 pathways, the receptor level crosstalk is insufficient for cadherin-11 to influence hMSC proliferation and an additional level of crosstalk is required between β-catenin and DUSP1. This has important implications for experiments concerning the control over hMSC proliferation.

Computational models are valid and useful in the range of biological systems they represent and for the biological data that is available to support them. In this study, we used an ODE model of the form described by Mendoza and Xenarios, 2006 to obtain a network that represents the qualitative characteristics of the RTK and cadherin-11 pathways and the crosstalk between them. This choice was made because we had a multitude of experimental data in the form of the relative abundance of active and inactive forms (i.e., phosphorylated and unphosphorylated forms) of proteins in the network and no kinetic information (i.e., binding–unbinding rates). Thus, a more classical mass action (or similar) type of model would have been difficult to construct and also to interpret, as the experiments were comparing relative protein abundances instead of absolute quantification. Nevertheless, with a network that focused on relative activities of proteins at the steady state, such as ours, we were able to conclude that a multi-level crosstalk between the two modeled pathways is needed to support the experimental observations. With the sensitivity analysis, we were able to show that the set of parameters we used was not unique to produce the results we obtained. This also indicated that more detailed information on the relative amounts of active/inactive proteins other than the RTKs, cadherin-11, and ERK could improve our choice of parameters and make the model more accurate in the future.

Similar to the cadherin-11–RTK interaction, other receptor couples are known to engage in crosstalk in other cell types (Andl and Rustgi, 2004; Chiasson-Mackenzie and McClatchey, 2018). For example, E-cadherin and epidermal growth factor receptor (EGFR) have been shown to interact (Ramírez Moreno and Bulgakova, 2022). Similarly, N-cadherin interacts with fibroblast growth factor receptor (FGFR) (Nguyen and Mège, 2016; Kon et al., 2019; Nguyen et al., 2019). We suggest a computational reconstruction of the signaling pathways of these receptor pairs, using a similar approach as we presented here, will help to isolate the pathways from other cellular signaling and to discover new interactions between the receptor pairs, other than what could be tested experimentally.

In summary, we have shown that a crosstalk between β-catenin (downstream to cadherin-11) and an ERK inhibitor protein (e.g. DUSP1) is needed for the experimentally shown effect of cadherin-11 on hMSC proliferation. By investigating the multi-level crosstalk between cadherin and PDGFRs computationally, this study contributes to an improved understanding of the effect of cell surface receptors on hMSC proliferation. A detailed description of how hMSC proliferation is controlled by a multitude of cell surface receptors will provide new avenues for cell fate control and regenerative medicine therapies.

## ACKNOWLEDGMENTS

This research was financially supported by the Gravitation Program “Materials Driven Regeneration”, funded by the Netherlands Organization for Scientific Research (024.003.013). The Virtual Cell is supported by NIH Grant R24 GM137787 from the National Institute for General Medical Sciences. We thank Kailas Honasoge for independently performing a reproducibility check of the model simulations. We would like to thank Bert Callens for the preliminary data that led to this manuscript.

## DATA AVAILABILITY STATEMENT

The model created for this project and simulation data that support the findings of this study will be publicly available in the VCell model database once the paper is peer-reviewed. During the preprint period, all the data is available upon request.

