## Supplemental for "COMPUTATIONAL EVIDENCE FOR MULTI-LAYER CROSSTALK BETWEEN THE CADHERIN-11 AND PDGFR PATHWAYS"

### Supplementary material

#### Tables

**Table S1** Parameter sensitivity analysis for β-catenin initial activity ($Bcat$) versus the inhibition of DUSP1 by β-catenin ($\beta_{dusp1_{by_{Bcat}}}$). Steady state activity levels of CyclinD1 and Erk are given for each parameter set in the search space.

| Set A | Bcat | $\beta_{\mathrm{dusp1}_{\mathrm{by}_{\mathrm{Bcat}}}}$ | cyclinD1 | erk |
|---|---|---|---|---|
| 0 | 0.1 | 0.1 | 0.5 | 0.4 |
| 1 | 0.1 | 0.5 | 0.5 | 0.4 |
| 2 | 0.1 | 1 | 0.5 | 0.5 |
| 3 | 0.1 | 5 | 0.3 | 0.6 |
| 4 | 0.1 | 7 | 0.2 | 0.7 |
| 5 | 0.1 | 10 | 0.2 | 0.7 |
| 6 | 0.3 | 0.1 | 0.5 | 0.4 |
| 7 | 0.3 | 0.5 | 0.5 | 0.4 |
| 8 | 0.3 | 1 | 0.5 | 0.5 |
| 9 | 0.3 | 5 | 0.3 | 0.6 |
| 10 | 0.3 | 7 | 0.2 | 0.7 |
| 11 | 0.3 | 10 | 0.2 | 0.7 |
| 12 | 0.5 | 0.1 | 0.5 | 0.4 |
| 13 | 0.5 | 0.5 | 0.5 | 0.4 |
| 14 | 0.5 | 1 | 0.5 | 0.5 |
| 15 | 0.5 | 5 | 0.3 | 0.6 |
| 16 | 0.5 | 7 | 0.2 | 0.7 |
| 17 | 0.5 | 10 | 0.2 | 0.7 |
| 18 | 0.7 | 0.1 | 0.5 | 0.4 |
| 19 | 0.7 | 0.5 | 0.5 | 0.4 |
| 20 | 0.7 | 1 | 0.5 | 0.5 |
| 21 | 0.7 | 5 | 0.3 | 0.6 |
| 22 | 0.7 | 7 | 0.2 | 0.7 |
| 23 | 0.7 | 10 | 0.2 | 0.7 |
| 24 | 1 | 0.1 | 0.5 | 0.4 |
| 25 | 1 | 0.5 | 0.5 | 0.4 |
| 26 | 1 | 1 | 0.5 | 0.5 |
| 27 | 1 | 5 | 0.3 | 0.6 |
| 28 | 1 | 7 | 0.2 | 0.7 |
| 29 | 1 | 10 | 0.2 | 0.7 |

**Table S2**

Parameter sensitivity analysis for activation of DUSP1 by ERK versus inhibition of DUSP1 by β-catenin ($a_{dusp1_{by_{erk}}}$ and $\beta_{dusp1_{by_{Bcat}}}$). Steady state activity levels of cyclinD1 and Erk are given for each parameter set in the search space.

| Set | $\mathrm{a}_{\mathrm{dusp1}_{\mathrm{by}_{\mathrm{erk}}}}$ | $\beta_{\mathrm{dusp1}_{\mathrm{by}_{\mathrm{Bcat}}}}$ | cyclinD1 | erk |
|---|---|---|---|---|
| 0 | 0.1 | 0.1 | 0.5 | 0.5 |
| 1 | 0.1 | 0.5 | 0.4 | 0.5 |
| 2 | 0.1 | 1 | 0.4 | 0.5 |
| 3 | 0.1 | 5 | 0.3 | 0.6 |
| 4 | 0.1 | 10 | 0.2 | 0.7 |
| 5 | 0.5 | 0.1 | 0.5 | 0.4 |
| 6 | 0.5 | 0.5 | 0.5 | 0.5 |
| 7 | 0.5 | 1 | 0.4 | 0.5 |
| 8 | 0.5 | 5 | 0.3 | 0.6 |
| 9 | 0.5 | 10 | 0.2 | 0.7 |
| 10 | 1 | 0.1 | 0.5 | 0.4 |
| 11 | 1 | 0.5 | 0.5 | 0.4 |
| 12 | 1 | 1 | 0.5 | 0.5 |
| 13 | 1 | 5 | 0.3 | 0.6 |
| 14 | 1 | 10 | 0.2 | 0.7 |
| 15 | 5 | 0.1 | 0.6 | 0.3 |
| 16 | 5 | 0.5 | 0.6 | 0.4 |
| 17 | 5 | 1 | 0.6 | 0.4 |
| 18 | 5 | 5 | 0.3 | 0.6 |
| 19 | 5 | 10 | 0.2 | 0.7 |
| 20 | 10 | 0.1 | 0.7 | 0.3 |
| 21 | 10 | 0.5 | 0.6 | 0.3 |
| 22 | 10 | 1 | 0.6 | 0.4 |
| 23 | 10 | 5 | 0.4 | 0.6 |
| 24 | 10 | 10 | 0.2 | 0.7 |

**Table S3**

Parameter sensitivity analysis for activation of ERK by MEK versus inhibition of ERK by DUSP1 ($a_{erk_{by_{mek}}}$ and $\beta_{erk_{by_{dusp1}}}$). Steady state activity levels of cyclinD1 and ERK are given for each parameter set in the search space.

| Set | $\mathrm{a}_{\mathrm{erk}_{\mathrm{by}_{\mathrm{mek}}}}$ | $\beta_{\mathrm{erk}_{\mathrm{by}_{\mathrm{dusp1}}}}$ | cyclinD1 | erk |
|---|---|---|---|---|
| 0 | 0.1 | 0.1 | 0.5 | 0.5 |
| 1 | 0.1 | 0.5 | 0.5 | 0.5 |
| 2 | 0.1 | 1 | 0.4 | 0.4 |
| 3 | 0.1 | 5 | 0.4 | 0.3 |
| 4 | 0.1 | 10 | 0.3 | 0.2 |

| | | | | |
|---|---|---|---|---|
| 5 | 0.5 | 0.1 | 0.5 | 0.5 |
| 6 | 0.5 | 0.5 | 0.5 | 0.5 |
| 7 | 0.5 | 1 | 0.5 | 0.5 |
| 8 | 0.5 | 5 | 0.4 | 0.3 |
| 9 | 0.5 | 10 | 0.3 | 0.2 |
| 10 | 1 | 0.1 | 0.5 | 0.6 |
| 11 | 1 | 0.5 | 0.5 | 0.5 |
| 12 | 1 | 1 | 0.5 | 0.5 |
| 13 | 1 | 5 | 0.4 | 0.3 |
| 14 | 1 | 10 | 0.3 | 0.2 |
| 15 | 5 | 0.1 | 0.5 | 0.6 |
| 16 | 5 | 0.5 | 0.5 | 0.5 |
| 17 | 5 | 1 | 0.5 | 0.5 |
| 18 | 5 | 5 | 0.4 | 0.3 |
| 19 | 5 | 10 | 0.3 | 0.3 |
| 20 | 10 | 0.1 | 0.5 | 0.6 |
| 21 | 10 | 0.5 | 0.5 | 0.6 |
| 22 | 10 | 1 | 0.5 | 0.5 |
| 23 | 10 | 5 | 0.4 | 0.3 |
| 24 | 10 | 10 | 0.3 | 0.3 |

**Table S4**

Parameter sensitivity analysis for activation of cyclin-D1 by DUSP1 versus inhibition of ERK by DUSP1 ($a_{cyclind1_{by_{mek}}}$ and $\beta_{erk_{by_{dusp1}}}$). Steady state activity levels of cyclinD1 and ERK are given for each parameter set in the search space.

| Set | $a_{cyclinD1_{by_{dusp1}}}$ | $\beta_{erk_{by_{dusp1}}}$ | cyclinD1 | erk |
|---|---|---|---|---|
| 0 | 0.1 | 0.1 | 0.4 | 0.6 |
| 1 | 0.1 | 0.5 | 0.3 | 0.5 |
| 2 | 0.1 | 1 | 0.3 | 0.5 |
| 3 | 0.1 | 5 | 0.2 | 0.3 |
| 4 | 0.1 | 10 | 0.2 | 0.2 |
| 5 | 0.5 | 0.1 | 0.4 | 0.6 |
| 6 | 0.5 | 0.5 | 0.4 | 0.5 |
| 7 | 0.5 | 1 | 0.4 | 0.5 |
| 8 | 0.5 | 5 | 0.3 | 0.3 |
| 9 | 0.5 | 10 | 0.3 | 0.2 |
| 10 | 1 | 0.1 | 0.5 | 0.6 |
| 11 | 1 | 0.5 | 0.5 | 0.5 |
| 12 | 1 | 1 | 0.5 | 0.5 |
| 13 | 1 | 5 | 0.4 | 0.3 |
| 14 | 1 | 10 | 0.3 | 0.2 |
| 15 | 5 | 0.1 | 0.8 | 0.6 |

| | | | | |
|---|---|---|---|---|
| 16 | 5 | 0.5 | 0.7 | 0.5 |
| 17 | 5 | 1 | 0.7 | 0.5 |
| 18 | 5 | 5 | 0.6 | 0.3 |
| 19 | 5 | 10 | 0.6 | 0.2 |
| 20 | 10 | 0.1 | 0.9 | 0.6 |
| 21 | 10 | 0.5 | 0.8 | 0.5 |
| 22 | 10 | 1 | 0.8 | 0.5 |
| 23 | 10 | 5 | 0.8 | 0.3 |
| 24 | 10 | 10 | 0.7 | 0.2 |

**Figures**

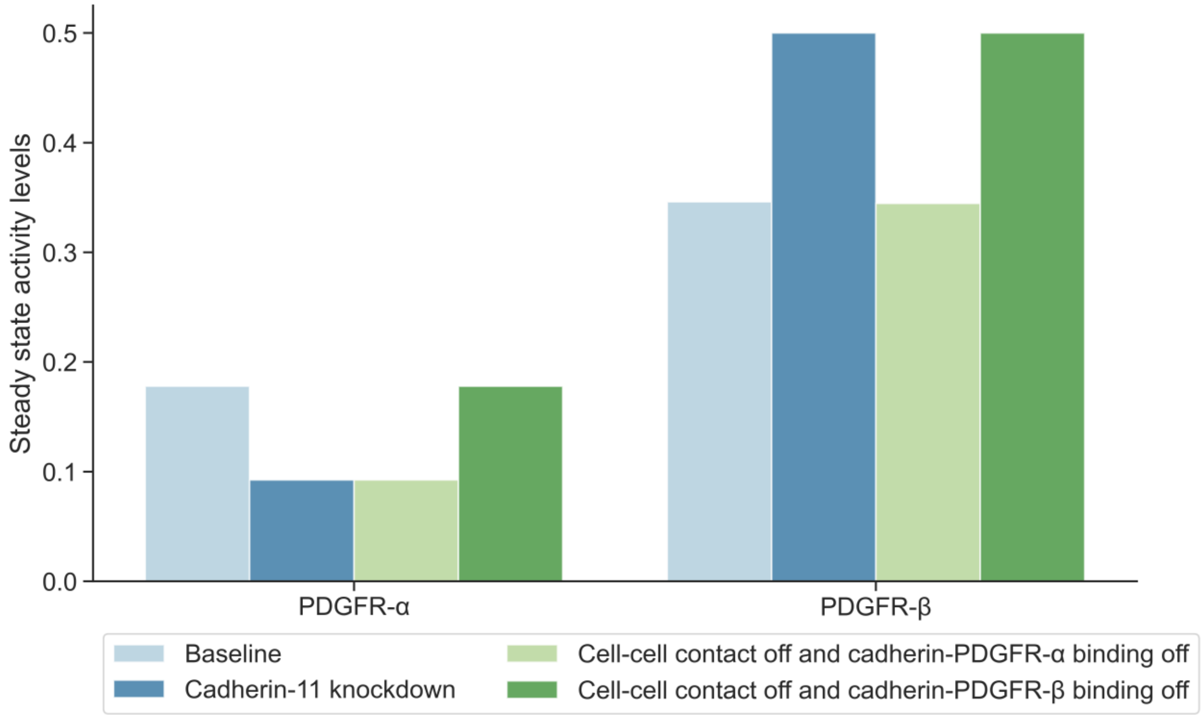


**Figure S1: Model parameters concerning the activity levels of PDGFR-α and PDGFR-β were adjusted to match the previous experimental studies (Madarampalli et al., 2019) and (Passanha et al., 2022)**. In the baseline model, PDGFR-β activity at the steady state is higher than PDGFR-α activity, in accordance with (Passanha et al., 2022). Cadherin-11 knockdown simulation resulted in lower than normal PDGFR-α and greater than normal PDGFR-β activity, in accordance with (Passanha et al., 2022). When cell–cell contact and cadherin-11 binding to PDGFR-α were disabled, lower than normal PDGFR-α activity but no change in the PDGFR-β activity were observed compared to the baseline model. When cell–cell contact and cadherin-11 binding to PDGFR-β were disabled, no change in the PDGFR-α activity level but an increase in PDGFR-β activity were observed compared to the baseline model.

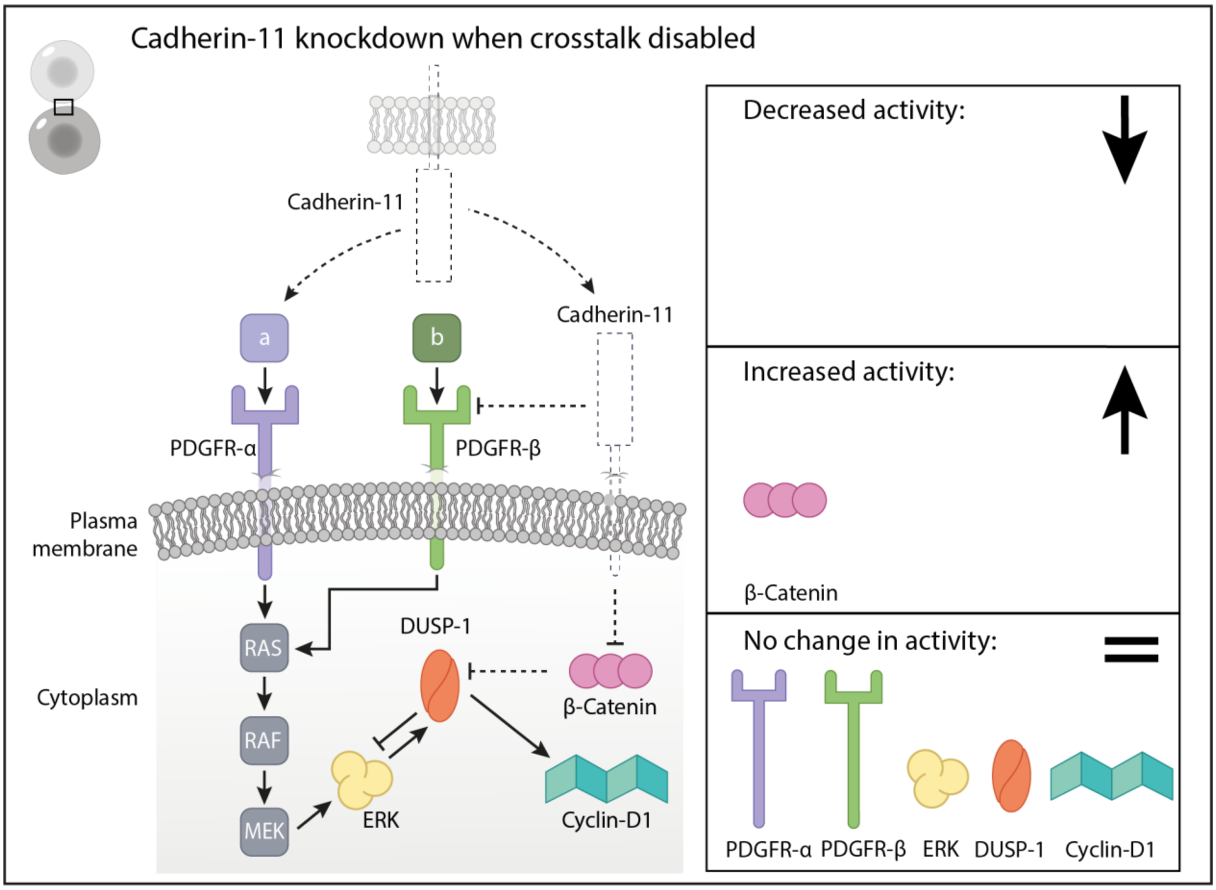


**Figure S2**: **Cadherin 11 knockdown simulation results when the proposed crosstalk is disabled:** The dashed lines indicate where the model components were modified in the simulation setup. No change was observed in the activity levels of growth factor receptors and proliferation-related signaling molecules in case of a cadherin-11 knockdown when the proposed crosstalk mechanisms were disabled.
